# Open Source Brain: a collaborative resource for visualizing, analyzing, simulating and developing standardized models of neurons and circuits

**DOI:** 10.1101/229484

**Authors:** Padraig Gleeson, Matteo Cantarelli, Boris Marin, Adrian Quintana, Matt Earnshaw, Eugenio Piasini, Justas Birgiolas, Robert C. Cannon, N. Alex Cayco-Gajic, Sharon Crook, Andrew P. Davison, Salvador Dura-Bernal, András Ecker, Michael L. Hines, Giovanni Idili, Stephen Larson, William W. Lytton, Amitava Majumdar, Robert A. McDougal, Subhashini Sivagnanam, Sergio Solinas, Rokas Stanislovas, Sacha J. van Albada, Werner van Geit, R. Angus Silver

## Abstract

Computational models are powerful tools for investigating brain function in health and disease. However, biologically detailed neuronal and circuit models are complex and implemented in a range of specialized languages, making them inaccessible and opaque to many neuroscientists. This has limited critical evaluation of models by the scientific community and impeded their refinement and widespread adoption. To address this, we have combined advances in standardizing models, open source software development and web technologies to develop Open Source Brain, a platform for visualizing, simulating, disseminating and collaboratively developing standardized models of neurons and circuits from a range of brain regions. Model structure and parameters can be visualized and their dynamical properties explored through browser-controlled simulations, without writing code. Open Source Brain makes neural models transparent and accessible and facilitates testing, critical evaluation and refinement, thereby helping to improve the accuracy and reproducibility of models, and their dissemination to the wider community.

## Introduction

Computational modeling is a powerful approach for investigating and understanding information processing in neural systems^1–3^. Data-driven models have played a central role in elucidating the mechanisms underlying synaptic transmission^4^, the action potential^5^ and dendritic integration^6^, and have been used more recently to investigate circuit function^7–12^. Such approaches are likely to become increasingly powerful as the amount of information available on the structure and dynamics of specific circuits increases, as a result of advances in connectomics^13, 14^, functional activity mapping^15^ and multi-cell and robotic patch clamping^16, 17^, together with datasets from large-scale brain initiatives^18–20^. Developing more accurate, data-driven computer simulations of neurons and circuits^7–9,21^ is increasingly important for integrating such multi-scale datasets into physico-chemically consistent models, and for helping to identify the key mechanisms involved in information processing and storage in health and disease.

Incorporation of newly discovered anatomical and functional properties of circuits has enabled the construction of well-constrained models of increasing biological detail ^7,9, 10^. However, developing data-driven models of neurons and circuits from a raw dataset takes a considerable amount of time and skill despite well established tools^22–25^. Additionally, as the biological realism of such models grow, more complex and lengthy machine-readable descriptions are required that depend on specialized tools including domain-specific simulators and numerical libraries. Running large-scale circuit models can also require high performance computing (HPC) facilities, which brings an additional layer of complexity to setting up simulations and managing the resulting data set. To scrutinize, systematically modify and reuse such biological detailed models, considerable specialist knowledge is required to structure the large code bases, databases and ancillary tools. These technical barriers are of concern as they prevent access to the structural and functional properties of models by the wider neuroscience community. This lack of accessibility and transparency limits the scientific scrutiny of models, impeding their refinement and reproducibility as well as their dissemination and adoption.

The extent and complexity of biologically detailed models also poses the technical challenge of ensuring their code bases are robust, reliable and error free. Common errors in the software implementation of such models include typos in equation definitions and parameter values, unit conversions, inconsistent use of temperature dependencies and disconnected neuronal morphologies. In addition, many existing neuronal models are described in a single monolithic block of code making them difficult to understand, test and integrate in larger scale circuit simulations. In contrast, modular programs with standardized building blocks can be more easily tested with automated routines and then assembled into larger structures, minimizing errors. In neuroscience, standardized model descriptions, such as NeuroML^26,27^, provide a defined structure that could be used for such a modular approach. Moreover, large-scale model development is beginning to involve groups of developers^28^ using version control software such as Git^29^, practices that are common in industry, but remain the exception rather than the rule in academia^30^. Developing, refining and maintaining ever more complex models of neural systems that are accurate, reliable and robust, therefore requires greater adoption of strategies currently used in open source software engineering for developing, managing, testing and validating large-scale code bases.

To address these challenges we have developed Open Source Brain (OSB; www.opensourcebrain.org), an open source web-based resource for visualizing, simulating, disseminating and collaboratively developing standardized models of neurons and circuits. OSB hosts a range of published neuronal and circuit models from multiple brain regions including the neocortex, cerebellum and hippocampus. The morphology and internal structure of models on OSB is automatically visualized in a form familiar to neuroscientists. Moreover, their functional properties can be explored by simulating models through the browser without installing programs or writing code. This is achieved by utilizing tools and best practices from the open source software development community, harnessing web browser and customized middleware technologies and integrating them with recently developed standardized model descriptions^26,27,31^. Unlike previous repositories, OSB provides infrastructure for collaboratively developing, refining and automatically testing models, enabling them to evolve as new information becomes available. By making models accessible and transparent, encouraging scientific evaluation and enabling further model development through collaboration, OSB provides an online resource of well-tested standardized models for use by the wider neuroscience community.

## Results

### The Open Source Brain Resource

Open Source Brain consists of a web-based content management system with a hub and spoke structure for linking open source repositories containing standardized models of neurons and circuits to users and developers. OSB provides powerful tools to visualize, analyze and simulate models through standard web browsers (**Fig. 1a**). These features, which expose the properties of models in a transparent manner, were made possible by defining models in the structured model specification languages NeuroML^26,27^ or PyNN^31^, since these can be parsed by specialized software to extract and visualize key aspects of the model (e.g. biophysical parameters, cell morphology) and transform them into a format that is familiar to neuroscientists. They also contain all the information required to generate scripts for simulations (**Fig. 1b**). OSB additionally facilitates access to metadata associated with the models, including their provenance, and has links to wikis and issue trackers, allowing users to discuss their properties, performance and any technical issues they may have. Models on OSB are hosted on public version control repositories (e.g. GitHub^29^), which provide functionality to track and manage changes to the source code. Tools are provided for developers to assist in the conversion of models developed in other languages (e.g. NEURON^24^, NEST^22^ and MATLAB) into standardized NeuroML and PyNN formats, together with infrastructure for automatically testing model validity (**Fig. 1a,b**). This combination of open source infrastructure and model standardization enables the front end of OSB to deliver models in familiar formats that can be understood and used by the wider neuroscience community, while the deep links to GitHub on the backend enable the software infrastructure for flexible collaborative model development and testing. The main web interface of OSB is at http://www.opensourcebrain.org.

**Figure 1.**
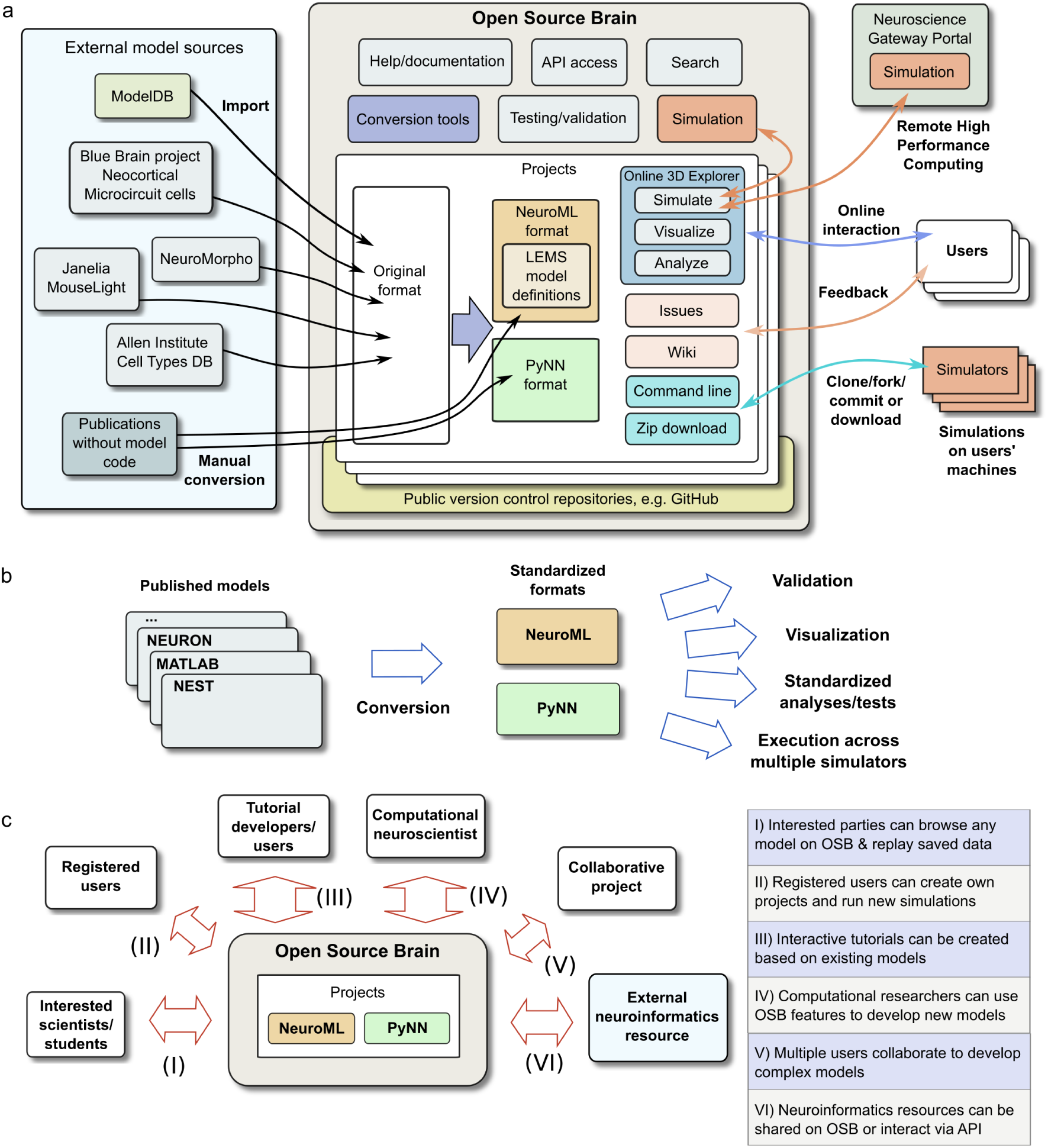
Overview of the Open Source Brain repository. **a**, Functionality of the Open Source Brain repository (OSB) and interactions with users and external resources. Left: original sources of models for projects on OSB. Center: OSB resources for helping to convert the models to standardized formats, to validate compliance to the standards and to test the model code, together with functionality for automatically visualizing, analyzing and simulating models in standardized formats through a web browser. A search function is provided for locating projects, together with an application programming interface (API) to access OSB functionality. Right: user interaction with individual projects for visualizing, analyzing and running simulations of the models can either be through the OSB graphical interface or through the command line. Wikis enable feedback and issues can be opened. Project code can be can be cloned/forked/committed using standard open source workflows or downloaded as zipped releases. Simulations can be set running locally on the OSB server or can be submitted to the Neuroscience Gateway Portal for execution on their supercomputing facilities. **b**, Depiction of increased functionality following converting published models from simulator specific formats into standardized representations. This includes automatic validation, visualization, analysis and simulations on different platforms, using a variety of generic tools. **c**, Usage scenarios for Open Source Brain projects containing standardized models in NeuroML or PyNN, depending on users’ goals and level of computational expertise.

Visualization of models and browser-controlled simulations on OSB rely on structured descriptions of models that are independent of the simulator or numerical integration methods used, together with a web-based graphical model interpreter. We use two languages for this framework: NeuroML^26,27^ and PyNN^31^. NeuroML is a widely used standard for declarative model description in neuroscience. NeuroML version 2 (the version we refer to in this paper) is built on a flexible low-level language, LEMS (low entropy model specification), that enables a wide range of physico-chemical processes to be defined^26^. The NeuroML2/LEMS framework allows diverse membrane conductances (Hodgkin-Huxley^5^, kinetic scheme, or any other formalism based on ordinary differential equations and state-dependent events), synaptic models (including chemical with short and long-term plasticity, electrical or analog/graded), point neurons or detailed neuronal morphologies and circuit connectivity to be specified and mapped to supported simulators (**Table 1**). PyNN is a Python API that allows scripts for network models of spiking neurons to be reused across multiple simulators, including NEURON^24^, NEST^22^, Brian^23^ and neuromorphic hardware. While PyNN and NeuroML have different approaches to model specification, these are interoperable: networks can be created with PyNN scripts and the structure exported to NeuroML format; cell models in NeuroML can be used in PyNN scripts and run on supported simulators (Methods). An advantage of having cell models from diverse sources in a common format is that analysis of the properties and behavior of the cells is greatly simplified. **Supp. Fig. 1** shows how the cell and channel properties of models developed originally in multiple formats can all be analyzed using the same procedural protocols (scripts) once they are in NeuroML.

**Table 1.** Models on Open Source Brain. A brief description of the properties of neuronal systems being modeled and the key model implementation features. Colors in first column correspond to the color code for brain regions in **Fig. 2**.

| Reference | Physiological properties | Model implementation | Link |
| --- | --- | --- | --- |
| Allen Institute Cell Types DB (Hawrylycz et al., 2016) | Morphologically detailed and point neuron models based on electrophysiological recordings from visual cortex neurons | Multicompartmental and Generalized Linear Integrate and Fire (GLIF) neuron models | URL |
| Brunel (2000) | Spiking network illustrating balance between excitation and inhibition | Integrate and Fire neurons; abstract network model; implementations in PyNN, NeuroML and NEST | URL |
| Hay et al. (2011) | Layer 5 pyramidal cell model constrained by somatic and dendritic recordings | Detailed neuronal morphology, non uniform channel distributions | URL |
| Izhikevich (2003) | Spiking neuron model reproducing wide range of neuronal activity | 2 variable point neuron model | URL |
| Markram et al. (2015) | Cell models from Neocortical Microcircuit of Blue Brain Project | Multicompartmental cell models of multiple cortical classes each with unique complement of active conductances | URL |
| Pospischil et al. (2008) | HH based model for different classes of cortical and thalamic neurons | Single compartment model with 5 ion channels reproducing a range of neuronal spiking behaviors | URL |
| Potjans and Diesmann (2014) | Microcircuit model of sensory cortex with 8 populations across 4 layers | Current based I&F neurons; implementations in PyNN, NeuroML and NEST | URL |
| Dura-Bernal et al. (2017) | Model of mouse primary motor cortex (M1) | Point neurons connected in columnar network based on realistic connection density distributions | URL |
| Smith et al. (2013) | Layer 2/3 cell model used to investigate dendritic spikes | Multicompartmental cell model with AMPA-R/NMDA-R mediated synaptic inputs | URL |
| Traub et al. (2005) | Single column network model containing 14 cell populations from cortex and thalamus | Semi-realistic neuronal morphologies; cell region specific distributions of active conductances; chemical & electrical synapses | URL |
| Cayco-Gajic et al. (2017) | Cerebellar granule cell layer network | Integrate and Fire based model for granule cell and anatomically constrained connectivity | URL |
| Maex and De Schutter (1998) | Cerebellar granule cell layer network | Conductance based point neuron models for granule and Golgi cells | URL |
| Solinas et al. (2007) | Cerebellar Golgi cell model | Conductance based model with abstract morphology | URL |
| Vervaeke et al. (2010) | Electrically connected cerebellar Golgi cell network model | Detailed Golgi cell model; realistic cell density & gap junction connectivity properties | URL |
| Bezaire et al. (2016) | Full scale network model of CA1 region of hippocampus | Detailed and abstract models of 10 cell types, realistic firing properties and connectivity parameters | URL |
| Ferguson et al. (2013) | Parvalbumin-positive interneuron from CA1 | Models are a customized form of the Izhikevich cell model | URL |
| Migliore et al. (2005) | Pyramidal cell from CA1 region of hippocampus | Multicompartmental cell model | URL |
| Pinsky and Rinzel (1994) | Simplified model of CA3 pyramidal cell | 2 compartment model; can be simulated as a single set of ODEs | URL |
| Wang and Buzsáki (1996) | Hippocampal interneuronal network model exhibiting gamma oscillations | Conductance based point neuron model used in 100 cell all-to-all network model | URL |
| Migliore et al. (2014) | Large scale olfactory bulb network | 600 unique multicompartmental mitral cells models along with simplified granule cells | URL |
| Boyle and Cohen (2008) | Model of body wall muscle from C. elegans | Point neuron model with 3 active conductances and internal $Ca^{2+}$ buffer | URL |
| FitzHugh (1961) | Simplified form of Hodgkin Huxley model | 2 variable cell model | URL |
| Hodgkin and Huxley (1952) | Classic investigation of the ionic basis of the action potential | HH model cell with current and voltage clamp inputs; model can be explored with interactive tutorial | URL |
| Prinz et al. (2004) | Pyloric network of the lobster stomatogastric ganglion system | 3 conductance based point neuron models connected via analogue (graded) synapses | URL |
| NeuroMorpho.Org | Digitally reconstructed neurons across multiple species and brain regions | All cells from NeuroMorpho.Org can be downloaded in NeuroML2; representative examples on OSB | URL |
| Janelia MouseLight | Reconstructed neurons with axonal projections stretching across the whole mouse brain | Cells downloaded from MouseLight website can be converted to NeuroML2; examples on OSB | URL |

**Figure 2.**
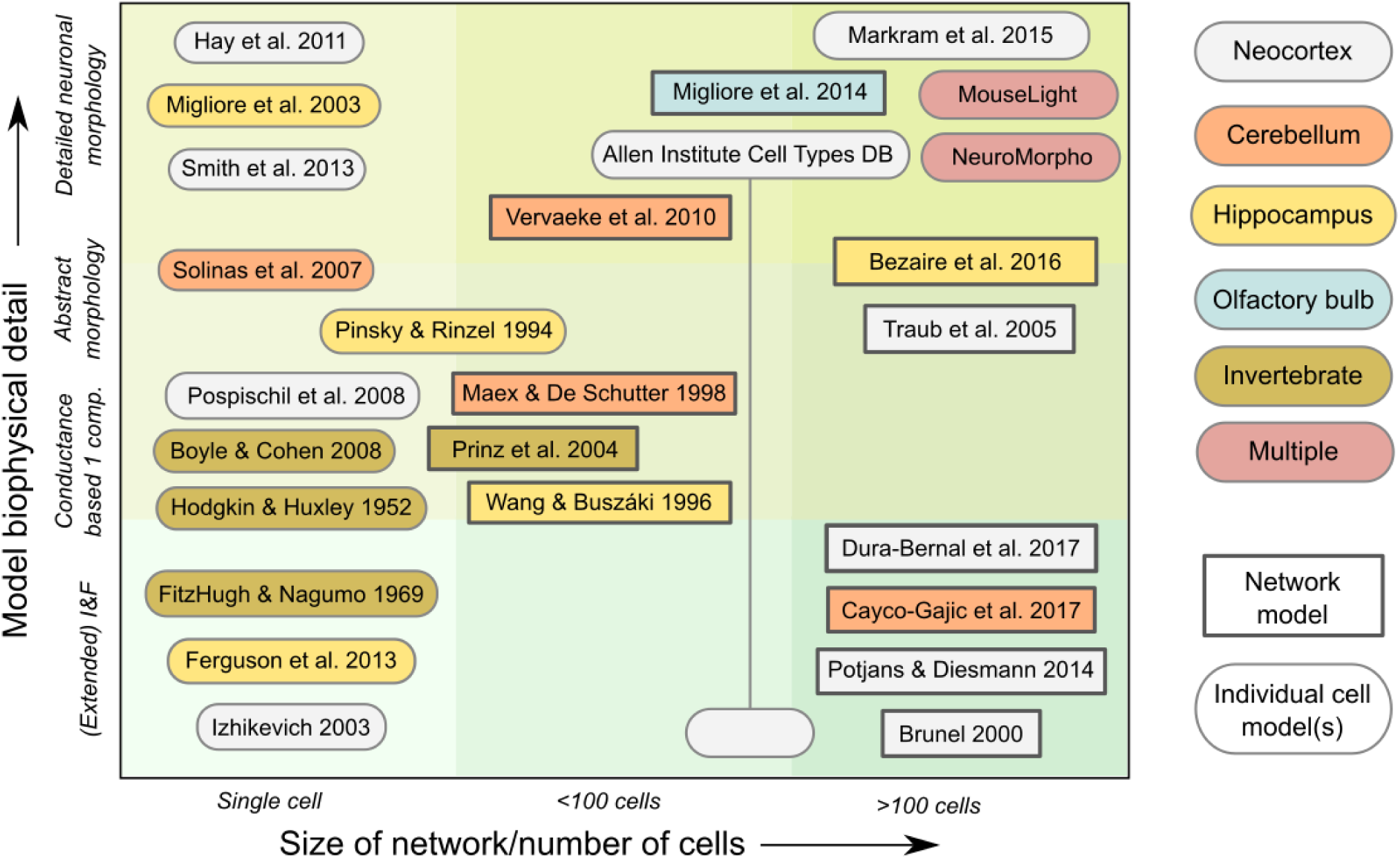
Standardized models on Open Source Brain span a wide range of scales and levels of biological detail. Graphical depiction of the NeuroML and PyNN based models on OSB, identified by author(s) of the paper in which model was originally described. Biophysical details range from point neuron models (e.g. Integrate and Fire (I&F)) to complex multi-compartment cell models based on detailed neuronal reconstructions. Some OSB projects contain individual cells while others contain multiple cell models, or network models with many instances of multiple cell types. Neuronal and circuit models cover a broad range of brain regions and include both vertebrates and invertebrates.

There are a number of different ways users can interact with models on OSB, depending on their goals and expertise in computational neuroscience (**Fig. 1c**). Scientists and students interested in learning about the properties of a model used in a scientific study can readily inspect model structure and parameters, and rerun previously recorded simulations, without registering as an OSB user. A more in-depth analysis of the model is possible following registration: users can run their own simulations to explore how parameters affect model behavior and can utilize graphical widgets to customize result visualization. When combined with the OSB framework for building tutorials, this can be used to develop online resources for teaching computational neuroscience to students and researchers. Another key OSB user group are scientists wishing to showcase/disseminate their models and to use the development and testing infrastructure of OSB in their model development. This group will initially be most interested in the tools provided to aid model conversion into a standardized format for use on OSB (**Fig.1a**). **Supp. Fig. 2** provides an overview of the steps required to add a model to OSB. Once converted, users will be able to use the tools for automated analysis and testing of models to help evaluate their accuracy and minimize software implementation errors. This supports the incremental evolution of a given model, ensuring that the intended behavior is not disrupted after each modification. Collaborative projects with multiple developers can also use OSB to build and test more complex models. Other neuroinformatics platforms can have their content shared on OSB or can pull details on all available models via the OSB web Application Programming Interface (API). For example there are deep links between OSB and ModelDB^32^, an archive of neuronal models in their original published formats.

### Standardized models currently on OSB

OSB currently hosts standardized curated models from multiple regions of the brain including neocortex^8,9,21,33–38^, cerebellum^10,39–41^, hippocampus^7,42–44^ and olfactory bulb^45^. These include single cell models from the Allen Institute Cell Types Database^19^ and the Blue Brain Project^8^, which have been converted to NeuroML, to enable visualization and simulation on OSB. A number of invertebrate and abstract network models have also been converted to standardized formats^5,46–48^. Standardized models that are currently available on OSB cover a wide range of biophysical detail and vary from single cells up to large-scale networks with hundreds or thousands of neurons (**Fig. 2**, **Table 1**; many other models on OSB are in development or in the process of conversion, besides those presented here). OSB is organized into projects containing sets of related cell and network models, and each project is linked to a single public version control repository which contains the model files.

### Visualization and analysis of model structure

Visualization of standardized models on OSB has been enabled through our contribution to the development of Geppetto (http://www.geppetto.org), an open source web platform built primarily in Java (server side) and JavaScript (client side) allowing models and data in various formats to be parsed, transformed and visualized through web browsers. The customized implementation of Geppetto for OSB provides extensive support for LEMS/NeuroML 2 models through the Java packages for NeuroML (jNeuroML) that we have developed (Methods). OSB searches for NeuroML files in the repositories associated with OSB projects, extracts structural information and metadata and allows visualization of them in 3D in the web browser in an interactive manner (**Fig. 3a**). All that is required for this functionality is WebGL (https://www.khronos.org/webgl) as is present in most modern browsers. Other properties of the model that can be extracted include information on the spatial dependence of parameters such as the density of ionic conductances on the plasma membrane. These can be presented in tabular form (in a “widget”), or as a pseudocolor density map superimposed on the neuronal morphology (**Fig. 3b**). Since models of ionic conductances are also specified in NeuroML format, the underlying mathematical expressions defining the rates of activation and inactivation can be extracted and plotted (**Fig. 3c**). Thus the types, distributions, densities and kinetic properties of the membrane conductances present on the model can be automatically exposed in a format familiar to electrophysiologists. Structured metadata in the NeuroML files is also extracted and presented through the web interface providing links to the source file and original data sources, thereby facilitating model provenance tracking.

**Figure 3.**
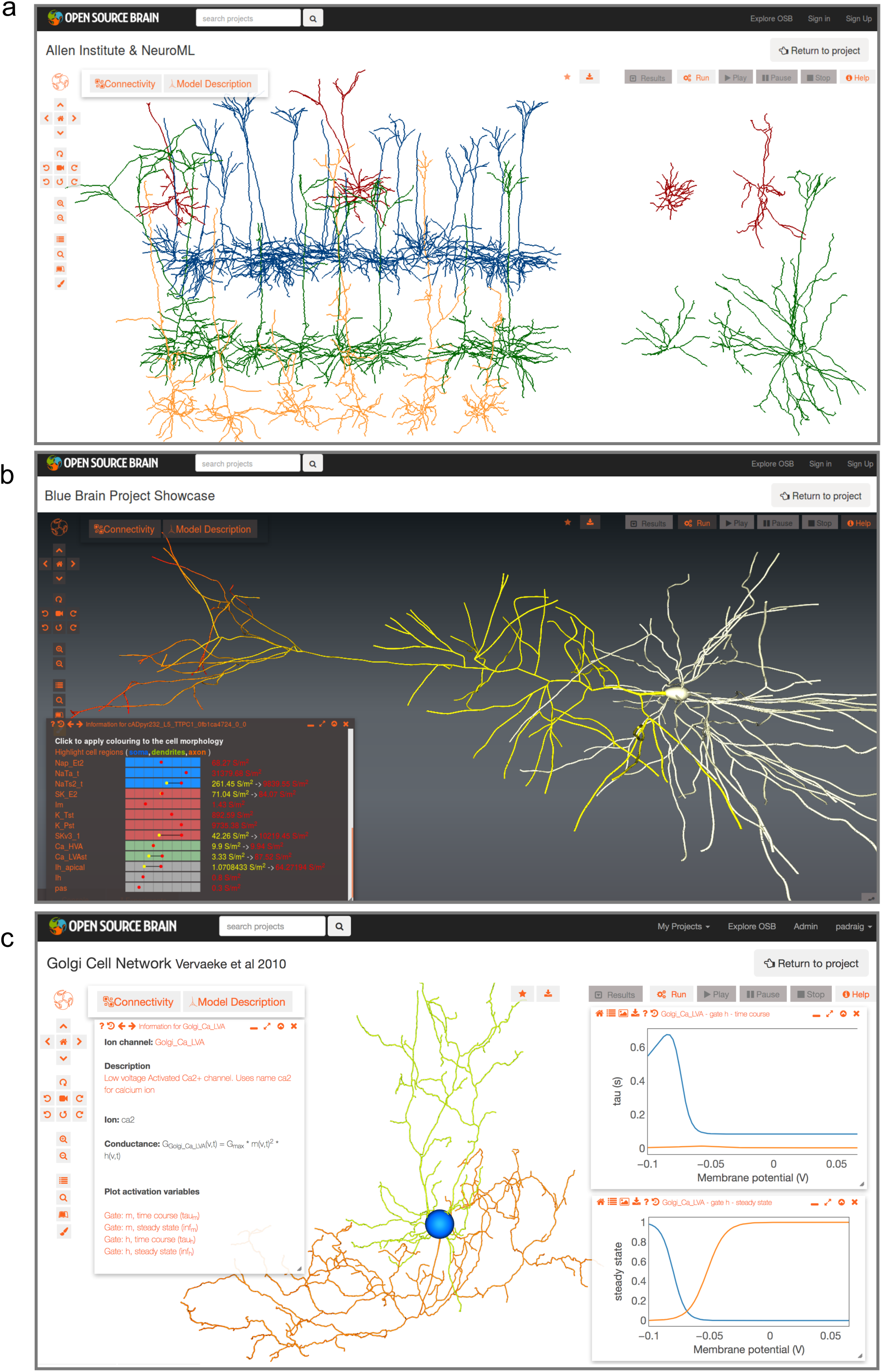
Visualization of models through the browser. **a**, Screenshot showing 37 cell models of visual cortex neurons from the Allen Cell Types Database (ACTD) visualized in a browser on the OSB website. These NeuroML conversions of models from ACTD have reconstructed morphologies with somatic spiking constrained by somatic electrophysiological recordings. Spiny (32 on left) and aspiny (5 on right) cells from Layers 2/3 (red), 4 (blue), 5 (green) and 6 (orange) are shown. **b**, Pyramidal cell from Layer 5 of the Blue Brain Project neocortical microcircuit model. Lower panel shows a widget summarizing the types and densities of ionic conductances present on the cell membrane. Individual conductances can be clicked and the regions of the cell containing that conductance will be highlighted. Main image of cell morphology demonstrates the non uniform distribution of Ih on the apical dendrite (low/yellow near the soma, high/red in the distal dendrites). **c**, Cerebellar Golgi cell model from Vervaeke et al. (2010). Cell regions have been highlighted (blue soma, green dendrites and orange axon). This multicompartmental cell model contains a number of conductances including a low voltage activated Ca2+ conductance (Ca LVA). Left panel shows widget-based description of the channel, including conductance expression and gating variables. Right panel shows voltage dependences of time constant (top) and steady state value (bottom) for the activation (orange, m) and inactivation (blue, h) gates for this conductance. Dendrite and axon diameters have been increased for clarity of figure presentation.

The 3D structure of circuit models is often complex, as it can include multiple neuronal layers, a range of cell types distributed at different densities and extensive interconnectivity, yet can exhibit highly specific synaptic connectivity between cell types. OSB facilitates network structure visualization by automatically generating multiple connectivity diagrams. This is possible because NeuroML descriptions of such networks contain lists of 3D locations of cells and the subcellular location of chemical and electrical synapses made between pre-and postsynaptic cells. **Fig. 4a** shows a single-column thalamocortical model consisting of multicompartmental neurons distributed over multiple cortical layers^21^. Here dendrites and somata are included but the synaptic connectivity is omitted for clarity. However, the synaptic connectivity of such circuits can be inspected in different ways using analysis widgets. A chord diagram (**Fig. 4b**) provides a convenient way to assess the density or sparsity of the synaptic connectivity. In contrast, the connectivity matrix (**Fig. 4c**) provides a more quantitative overview of the synaptic connections, showing the weights of connections between cells grouped by population. Lastly, the force-directed plot (**Fig. 4d**) provides a metric of the structure of the network. In a similar way, a cortical network consisting of point neurons^9^, which was specified in PyNN and exported to NeuroML, can be analyzed and compared with the more detailed model (**Fig. 4e-h**). For large-scale networks with a high level of biological detail, such as the recently developed CA1 circuit model^7^, OSB can progressively load parts of the network. For example, visualization of the gross structure of the circuit, distribution of cells and dendritic morphology does not require loading the synaptic connectivity matrix (**Fig. 4i**). However, this can be loaded in the background if required, enabling the properties of the synaptic connectivity to be visualized (**Fig. 4j-l**).

**Figure 4.**
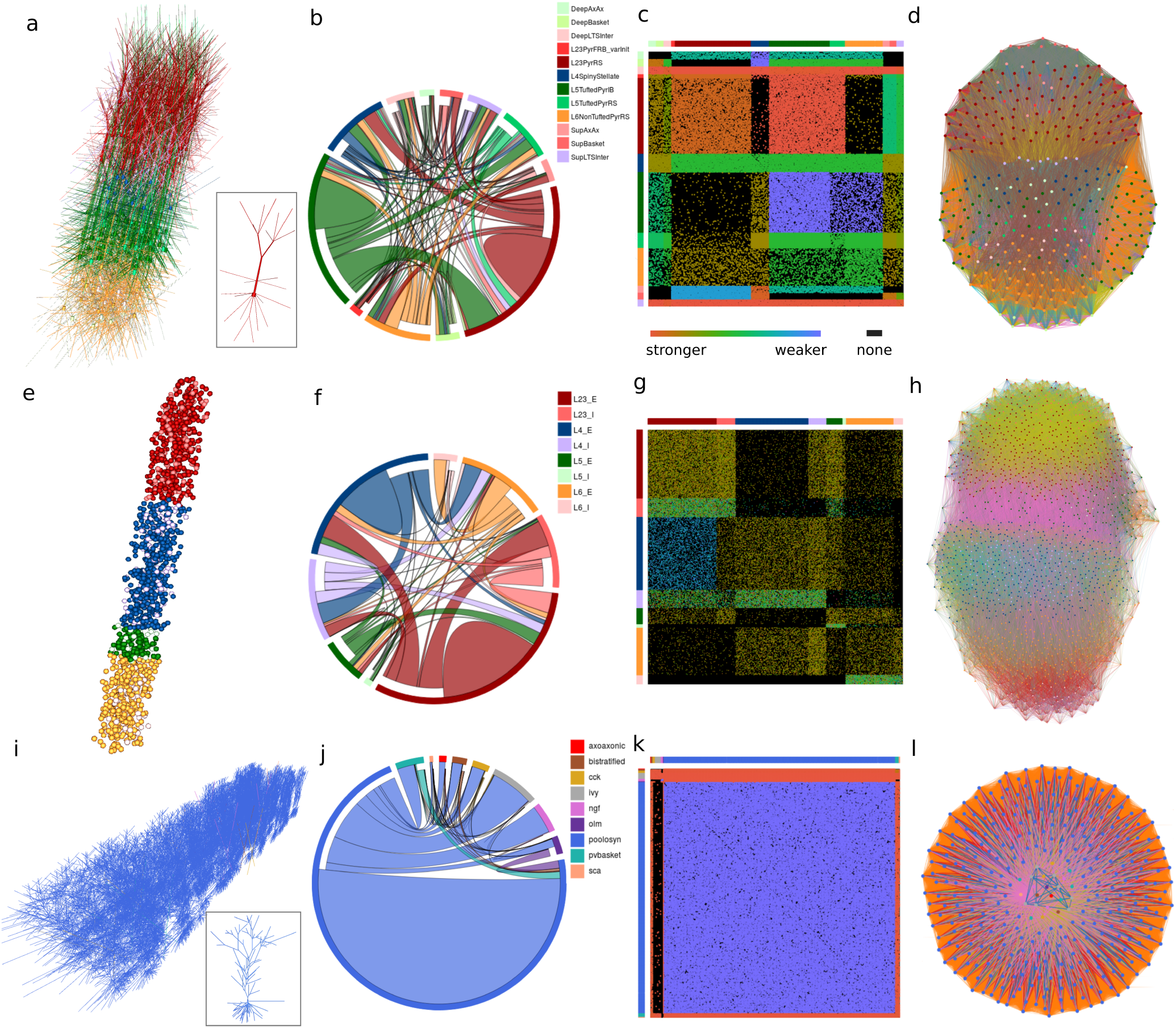
Analysis of network structure on Open Source Brain. **a**, Single-column thalamocortical network model from Traub et al. (2005), containing 336 cells (between 100 and 148 compartments each; 10% of full network) in 12 populations. Inset shows example pyramidal cell from layer 2/3. **b**, Chord diagram showing projections between populations (100K individual connections of 193 types). Outer ring color indicates population; the chords are attached to the presynaptic population outer ring segment, separated from the postsynaptic population ring segment and colored to match the postsynaptic population. Colors for populations match 3D view in (a). **c**, Adjacency matrix; lines on top and left indicate pre-and postsynaptic population colors respectively. Dots in matrix represent single connections, colored by weight. **d**, Force directed graph between circles representing each cell. Force, and so closeness, between pairs of cells is proportional to connectivity, resulting in grouping of highly connected cells. Clustering of L2/3, L4 and L6 (red, blue and orange respectively) can be seen. **e-h**, as for (a-d) but for point neuron spiking network model of Potjans and Diesmann (2014) with 1539 cells in 8 populations (E (excitatory) and I (inhibitory) from L2/3, L4, L5 and L6; 59.6K connections; 2% of full scale network). **i-l**, as for (a-d) but for network model of hippocampal CA1 region from Bezaire et al. (2016) with 311 pyramidal cells (example cell in inset of (i)) and 24 interneurons of different types. Network (1% of full scale) has over 1M connections, primarily between pyramidal cells. All images based on screenshots taken from interactive browser visualization of models on OSB.

### Visualization of functional properties through online simulation

Taking advantage of the simulator-independent representations of models on OSB, we extended the functionality of Geppetto to allow models to be run through the browser, without writing code. Instructions for simulating the model are fed to the backend, where the scripts are automatically generated and executed. Small-scale, short simulations can be run quickly on computing resources provided by OSB, and larger scale/longer computations can be submitted for execution through the Neuroscience Gateway at the San Diego Supercomputer Center^49^ (**Fig. 1a**). Exporting to NetPyNE format (a Python package building on NEURON, http://netpyne.org) allows parallel execution of models across hundreds of processors (see Methods). Upon completion, the data generated are sent back to the browser for visualization (**Fig. 5**,**6**).

**Figure 5.**
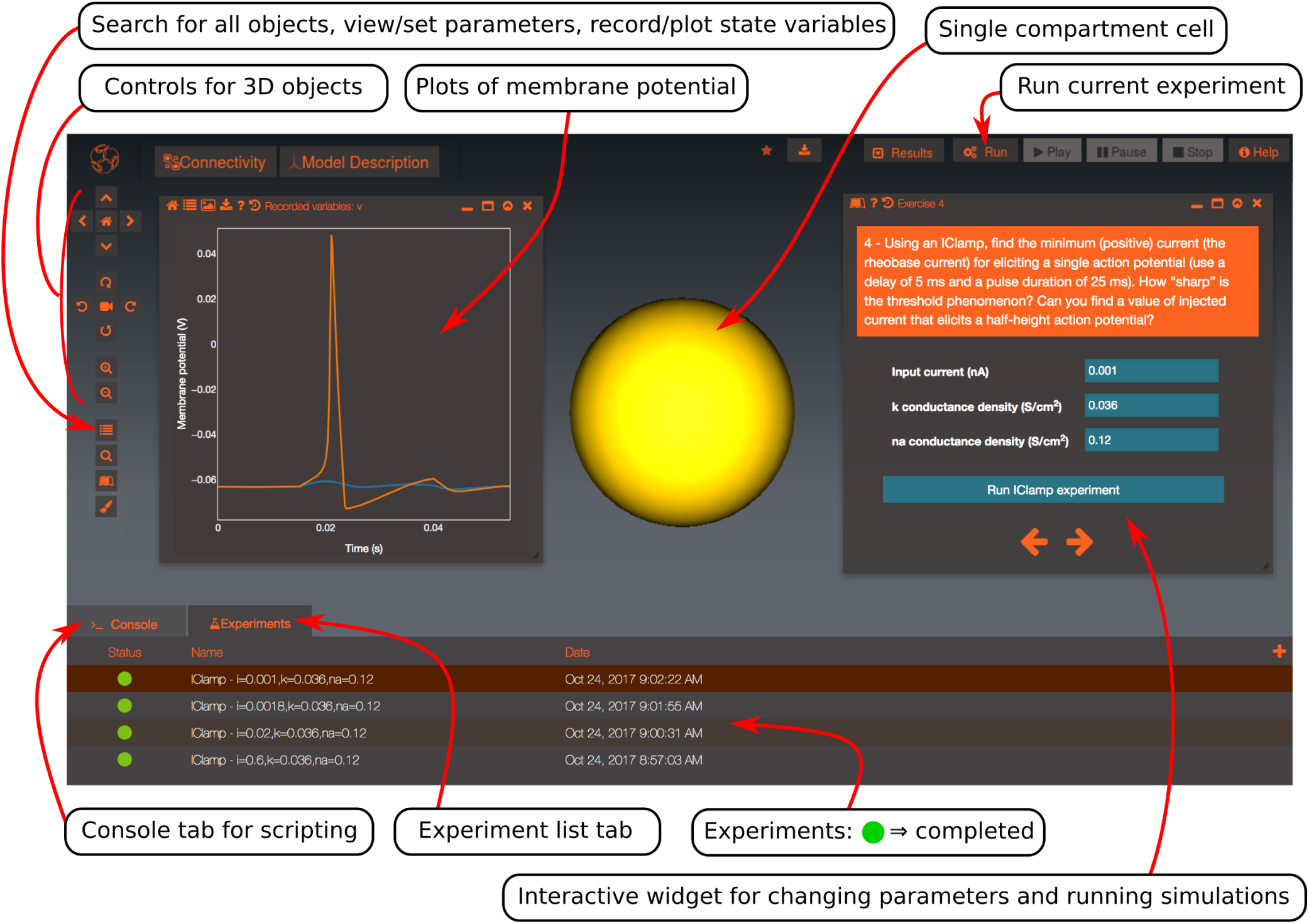
Interactive online tutorial on the action potential and control and execution of simulations. An annotated screenshot of an OSB project for a single compartment neuron with Hodgkin-Huxley model conductance (Hodgkin and Huxley, 1952), including graphical elements that have been added with the tutorial builder. The single compartment model cell (yellow, sphere), a tutorial widget for altering current, channel densities and running simulations (right), a list of previously run experiments with changed simulation parameters (bottom) and membrane potential plot showing spiking (orange) and subthreshold (blue) recordings (left) are shown. Controls for manually changing parameters, recording state variables and running simulations (as opposed to using the interactive tutorial widget) are also highlighted by the ‘Search for objects, set/view parameters, record/plot state variables’ arrow and the ‘Run current experiment’ arrow.

**Figure 6.**
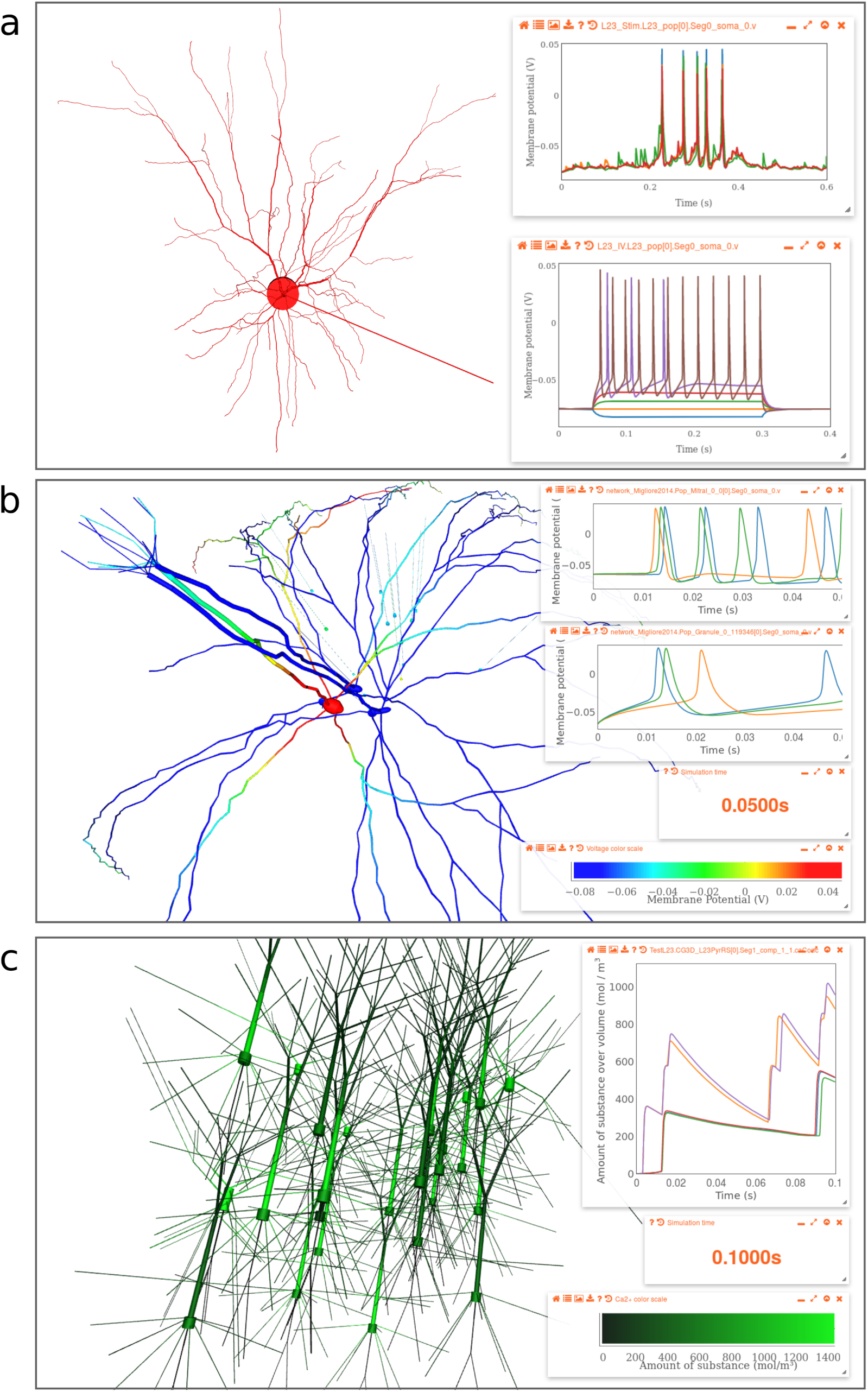
Visualization of simulations through the browser. **a**, Layer 2/3 pyramidal cell project from Smith et al. (2013). Left, cell morphology visualized on OSB red. Top panel on right: plot of membrane potential at 5 locations from an OSB-based simulation of this cell during background synaptic stimulation. Bottom panel: plot of firing frequency versus injected current for multiple OSB simulations. **b**, Small network with 3 mitral and 15 granule cells from Migliore et al. (2014). Plots on right show time course of somatic membrane potentials for mitral and granule cells. Widgets below show simulation time during replay and color scale for the membrane potentials at different locations, which are plotted dynamically on the cell morphologies. **c**, Left: A small network of Layer 2/3 pyramidal cells and interneurons from Traub et al. (2005). Right hand plot shows time course of intracellular calcium concentration during the simulation. Right bottom, scale for recorded calcium concentration overlaid on cell morphologies.

To enable the functional properties of models to be explored, model parameters (e.g. input currents, firing rates, cellular/synaptic properties) can be edited through the browser (**Fig. 5**). Running multiple simulations enables characteristic neuronal properties, such as the relationship between injected current and voltage, to be investigated (**Fig. 6a**). Differing levels of detail can be recorded in cells, from activity at only the soma, to the voltage in every compartment. Recorded data can be replayed as a time series or, if the model is spatially extended, a pseudocolor code can be used to indicate the voltage (**Fig. 6b**) or calcium concentration (**Fig. 6c**) across the morphology or across a population of neurons.

### Simulation management and tutorials

We have built a system for managing and storing simulations on OSB. This enables registered users to manage multiple simulations and to perform analyses on them (**Fig. 5**). In addition, the layout of the 3D canvas and associated widgets can be saved between sessions. This feature is supported by a command line console in the OSB interface that provides a history log of modifications to the model/graphical elements (see Methods). Interaction by users directly through the console can also speed up analysis and the generation of custom layouts. The simulations carried out on OSB can also be downloaded to the user’s computer or to automatically uploaded to Dropbox for offline analysis (see Methods).

The online nature of OSB, together with its simulation and management features make it well suited for demonstrating the principles of neurophysiology in an interactive and accessible format. To this end we have built a framework for constructing online tutorials that can be used to explain concepts through presentation of figures and simulations. These features enable interactive tutorials and virtual experiments to be constructed that can be used to teach basic concepts in neurophysiology and computational neuroscience without the barrier of having to write code or install specialist simulators. To illustrate this functionality we have extended a preexisting tutorial on the Hodgkin Huxley model of the action potential for use on OSB (**Fig. 5**).

### Conversion, development and testing of standardized models

Models of neural systems can be developed directly in standardized formats^10^, but most have been developed and defined in simulator-specific languages^32^. To utilize the full functionality of OSB, simulator-specific models need to be converted to standardized formats and deposited in an open source code repository that is linked to OSB. To facilitate this process we have developed a range of documentation and tools to facilitate model conversion and development to standardized formats (Methods). **Supp. Fig. 2** provides an overview of how these tools can be used at each stage of conversion of an existing model for use on OSB. Source code repositories for the models on OSB are generally hosted on GitHub, a code hosting platform that provides a range of resources for open source code development. These include version control software for recording changes and managing/merging different users’ branches/forks of the main repository, besides tracking issues and flagging stable releases.

Development of a model requires systematic testing of the code base. In software engineering, it is standard practice to run automated tests whenever a change is committed to a repository (“continuous integration”). To facilitate this on OSB we have developed the OSB Model Validation framework, which allows test configuration files to be added to a project. These specify the file to test, the simulator and the expected outcome of the simulation (e.g. times of spikes) with an appropriate tolerance. This approach is also useful in model conversion and refinement as it enables comparison of the original model behavior and the behavior of the standardized description. Tests can be run on a user’s local machine and with remote continuous integration services such as Travis-CI (Methods). The latter option ensures the full sequence of simulator installation, model execution and validation is run on every change to the model code, quickly highlighting deviation from expected behavior (and simulator compatibility issues). **Supplementary Table 1** shows the range of simulator specific tests on the OSB models discussed in this paper.

To facilitate further the distribution and testing of OSB models, we have used Docker (https://docker.com) to create self-contained software environments (images) with all simulator tools preconfigured, as well as verified stable releases of all models presented here (**Table 1**). Using these images, the full test suite can be run using a few commands on any operating system supporting Docker, currently comprising 324 individual tests across 21 simulator configurations in 26 projects (**Supp. Table 2**).

## Discussion

We have developed Open Source Brain, an open source web resource of standardized neuronal and circuit models and tools for collaborative model development. The core aims of this initiative are to make models of neural systems accessible and transparent to the wider neuroscience community, to encourage critical evaluation of model properties and behavior, to improve model validation and reproducibility and to facilitate collaboration in open source model development. To this end, the OSB platform enables web browser based visualization, analysis and simulation of models from single cells up to large-scale networks without the need to install software or write code. It also provides the infrastructure and tools for collaborative model development. This has been achieved by applying open source software development methodologies, utilizing standardized model descriptions, harnessing modern web technologies and building on community efforts by model and application developers to make their code open, usable and freely available. By making data-driven neuronal and circuit models more accessible, transparent and reproducible, and promoting collaborative development of standardized models, the OSB platform provides a powerful new resource for researchers and students investigating brain function in health and disease.

The distinctive features of OSB arise from the fusion of several recently maturing technologies. Automated model visualization and simulation stem from the use of structured models that are defined in simulator-independent standardized formats. Indeed, the use of the standardized languages, NeuroML2^26,27^ and PyNN^31^ are critical to the functionality of the platform. They have reached a level of maturity that allows a wide range of neuronal and circuit model types to be defined (**Fig. 2**, **Table 1**). Furthermore, their ability to describe basic physico-chemical processes confers the flexibility to define models of biochemical processes, as developed in systems biology^26^, suggesting that increasingly detailed models can be accommodated within these frameworks.

Visualization of the structural and functional properties of models relies on a host of web technologies including WebGL and JavaScript. The open source Geppetto platform makes use of these to allow users to interact with the underlying models in a graphical manner without requiring programming knowledge. To enable large-scale simulations, OSB is tightly integrated with the Neuroscience Gateway^49^ (NSG), which facilitates wider access to supercomputing facilities funded by the US National Science Foundation. OSB lowers the barriers for using these facilities for computational neuroscience, by enabling browser-controlled simulations of neurons and networks to be automatically submitted to NSG, providing large-scale parallel simulations across hundreds of processors (see Methods). Once simulations are completed, Geppetto provides the framework to retrieve the simulation data and to plot and manage it from the browser, thereby enabling accessible exploration of model behavior. Allowing a wider range of users access to large-scale simulations will enable critical evaluation of such complex models and provide a more informed discussion of their utility within the neuroscience community.

An important aspect of the workflow on OSB is the continuous open source development, refinement and testing of models. This is enabled through deep links to GitHub which provide the version control infrastructure for collaborative software development. Also key to developing robust reproducible models is the battery of new tools we have developed for automatically testing standardized model code. The OSB Model Validation framework, which utilizes the unit testing methodology from software engineering, ensures model behavior does not change when converted to standardized formats and when updates are made to the code. This helps maintain code quality and consistency, which is essential for developing complex large-scale models and when model components are used across labs and research projects.

The distinct functionality of OSB extends and complements that of ModelDB^32^, a well-established, extensive repository of models for computational neuroscience. ModelDB hosts model code in the original language in which it was developed and facilitates the reproduction of the results from their originating publications. It allows users to search based on model features, and provides information on the structure and properties of models that have been developed using the NEURON simulator. OSB builds on this functionality by focusing on hosting standardized models, which enable users to interact with and analyze models in greater detail (deep links allow users to find the same model on either platform). Moreover, OSB is designed to expose circuit-level properties including connectivity matrices and network dynamics. But perhaps more fundamental is that standardized models hosted on OSB are not frozen as they are in ModelDB, but can be developed, refined and improved using the infrastructure provided for open source software development, automated validation and testing of models. OSB actively encourages researchers to first submit their model to ModelDB following publication (**Supplementary Fig. 2**). The step of moving a model onto OSB is an indication that one or more parties (who may not be the original developers) wish to standardize and further develop the model, making it more accessible to the wider neuroscience community and extending it for use beyond the original publication.

OSB also interacts with other online resources that provide structured, annotated data that are invaluable for creating and validating neuronal models. For example, reconstructed neuronal morphologies from NeuroMorpho.Org and the Janelia MouseLight project (http://mouselight.janelia.org) can be automatically converted to NeuroML and visualized through OSB, and these morphologies can become the basis for new models when combined with the complement of membrane conductances for that cell type already in NeuroML. The Allen Institute Cell Types Database^19^ provides electrophysiological recordings and morphological reconstructions from cells in mouse visual cortex. Biophysically detailed cell models and point neuron models based on these data are present on OSB, as are cell models from the Blue Brain Project’s reconstruction of the microcircuitry of rat somatosensory cortex^8^, and a reduced version of a cortical column used in the Human Brain Project^20^ (HBP). Converting the neuronal models present in these networks to standardized formats also provides a valuable resource for developing new models of cortical circuits from modular, well tested building blocks.

OSB is designed so anyone can add a new project and use the functionality for developing testing and promoting their project, without the need for OSB administrators to get involved. This will enable the resource to grow in a way that is determined by the user’s interests. Models contributed by individual labs will be complemented with community developed models, including the HBP HippoCamp initiative, for which our CA1 network conversion (**Fig. 4i-l**) is a first major contribution, and the OpenWorm project^50^, which aims to create an in-silico model of the nematode *C. elegans*. In addition to contributing new models to the repository, academics can use the OSB online tutorial building functionality to build interactive teaching resources for undergraduates and postgraduates that combine text, theory and simulations. These can be used to explain the principles of the action potential, synaptic integration, dendritic processing and circuit function as well as how to build a computational model.

In the longer term, we intend to expand the functionality of the platform to include tighter integration with the experimental data used to build models and test their performance. Standardized data formats, such as Neurodata Without Borders (http://nwb.org), will be crucial for this. Direct comparison of experimental data with model properties will provide a new level of transparency and scientific evaluation for data-driven models. It is hoped that the software infrastructure presented here, together with these future developments, will lead to more accurate and predictive models of brain function in health and disease that can be understood and used by the wider neuroscience community.

## Acknowledgements

The OSB initiative was funded by the Wellcome Trust (086699, 101445, 095667). RAS is in receipt of a WT PRF (203048) and an ERC advanced grant (294667). In addition, the infrastructure to enable integration of OSB and the NSG was funded by the BBSRC-NSF/BIO program (BB/N005236/1 & NSF #1458840) and NSF #1458495. EP was supported on the EU Marie Curie Initial Training Network CEREBNET (FP7-ITN-PEOPLE-2008; 238686). SD-B and WWL were funded by NIBIB U01EB017695, DOH01-C32250GG-3450000 and NIH R01EB022903. MLH and RAM were funded by NIH grant DC009977 and MLH by NIH NS11613. BM was funded by Fundação de Amparo à Pesquisa do Estado de São Paulo (FAPESP), proc. 2017/04748-0. SC and JB were partially funded by NIH R01MH106674 and NIH R01EB021711. SJvA was funded by EU 7th Framework Program (FP7/2007-2013) under grant agreement n° 604102 (Human Brain Project, Ramp up phase) and EU Horizon 2020 research and innovation program under grant agreement n° 720270 (HBP, SGA1). The work of AE, JB and RS was partially funded by the Google Summer of Code program, coordinated through the INCF. We are grateful to Marianne Bezaire for assistance in converting the CA1 network model^7^, and Eilif Muller for help converting the Blue Brain Project cells^8^. We would also like to thank the other developers who have made contributions to the models on OSB and supporting applications. Specific contributions can be found on individual GitHub repositories.

## Author Contributions

PG and RAS conceived the Open Source Brain project. PG, RAS, SC, RCC, MLH, BM, JB, EP contributed to the NeuroML language and tools. PG and APD developed the PyNN/NeuroML toolchain. MC oversaw the development of Geppetto functionality and MC, GI, AQ, BM, ME, PG, RAS contributed to the design and development of the Geppetto features. MC, AQ, BM, ME, EP, PG, GI, RAS contributed to the design and development of the OSB platform, including the Redmine based frontend. PG, BM, EP, NAC-G, RS, AE, APD, JB, SS, SD-B, SJvA, WvG, SL converted models to the standardized formats. BM and PG developed the OMV testing framework. PG and RAM developed links between OSB and ModelDB. PG, AQ, AM, SS developed the interface with NSG. SD-B and WWL contributed to the NetPyNE mapping for models. PG and RAS wrote the manuscript with input from all authors.

## Competing Interests

The work carried out for this publication by Robert Cannon was conducted under contract from UCL to his employer, Annotate Software Limited in which he is also a shareholder. MetaCell Ltd. was also contracted by UCL to develop some of the features of the Open Source Brain software. MC, GI, SL declare financial interest in MetaCell Ltd. The other authors declare no competing financial interests.

## Methods

### OSB web frontend: project and user management

The main OSB web interface (at http://www.opensourcebrain.org) is based on a heavily customized version of Redmine (http://www.redmine.org). This platform, developed in Ruby on Rails (http://rubyonrails.org), allows users to create accounts, make new projects and add other users to them, create user groups, and associate a version control repository (hosted on GitHub (https://github.com), BitBucket (https://bitbucket.org), SourceForge (https://sourceforge.net) etc.) to each project. We have extended this framework with a new user interface, and added custom fields for projects with metadata relevant to the model, brain region, species and simulators supported. Close integration with platforms hosting the version control repositories allows the files and change history to be deeply integrated into the OSB interface. Issues raised on GitHub, or forks of the repository are highlighted on the OSB page. Wikis describing the installation/usage of the models can be added directly on OSB, or can retrieve and display content from README files in the associated repository. An API is provided to programmatically access information on all current projects and metadata (https://github.com/OpenSourceBrain/OSB_API). OSB projects’ repositories are automatically searched for NeuroML files (in either XML or HDF5 format) and these are presented to users for visualization in the integrated Geppetto application.

### OSB web frontend: model visualization & simulation

Geppetto (http://www.geppetto.org) is an open source web platform enabling parsing of models and experimental data on a central server and presenting these for visualisation and simulation in standard web browsers. The application was originally created in the OpenWorm project^50^, but it has developed into a modular platform with a number of parties contributing features. It was significantly extended for use in OSB with the ability to visualize neuronal models in 3D, to simulate them remotely and present the resultant data to users in the browser. It uses the Java based NeuroML libraries mentioned below to handle the reading and writing of NeuroML models and conversion to simulator formats. A number of JavaScript packages are used on the client side for 3D visualization (WebGL; three.js), user interaction (React; D3) and plotting (Plotly). In addition to NeuroML, Geppetto can interpret and display SWC format^51^ as well as 3D objects in OBJ format (http://www.martinreddy.net/gfx/3d/OBJ.spec).

Geppetto provides a canvas for displaying and interacting with 3D objects (**Figs. 3**, **5**, **6**) as well as a number of “widgets” for displaying textual information, interaction elements such as buttons and menu items, and plots. Connectivity information extracted from networks can also be displayed in custom widgets (**Fig. 4**). Interaction with the 3D objects and widgets can be solely through the graphical interface, but Geppetto also provides an integrated console for interactions with all these elements through JavaScript (enabled by clicking on the Console tab at the bottom of the 3D view (**Fig. 5**)).

### NeuroML 2 & LEMS libraries

To facilitate the use of NeuroML models on OSB, we have developed a number of libraries for processing files in this standardized model specification language. These features have been developed in Java and Python, two of the most commonly used languages in computational neuroscience. A focus of the work has been on making these features available through easy to install packages. jNeuroML (https://github.com/NeuroML/jNeuroML) is a single package which gives access to all features which have been implemented in Java^26^. These include: natively parsing and simulating models specified in LEMS (including point neuron cell models/networks in NeuroML); converting NeuroML models to simulator-specific code (for currently supported simulators see **Suppl. Table 1**); importing other structured formats to LEMS (particularly SBML^52^ models); validating NeuroML files, as well as performing basic tests for consistency; converting 3D models to SVG and PNG images.

libNeuroML (https://github.com/NeuralEnsemble/libNeuroML) is a Python package for reading, editing and writing NeuroML files^53^. pyNeuroML (https://github.com/NeuroML/pyNeuroML) is a Python package which builds on libNeuroML and bundles a copy of jNeuroML, allowing access to all of its functionality from Python scripts (most importantly converting NeuroML models to simulator code, running them and reloading the results). Additionally, it has utility scripts for analysing channel properties (*pynml-channelanalysis* for NeuroML channels; *pynml-modchananalysis* for NEURON channels) and generating high resolution images and movies using POV-Ray (*pynml-povray*, http://povray.org). Both jNeuroML and pyNeuroML can be used as command line applications (*jnml* and *pynml* respectively) or as libraries to give access to these features in other Java or Python applications (e.g. jNeuroML is bundled with neuroConstruct^54^).

Both the Java and Python libraries can serialize NeuroML models as either XML or in binary HDF5 format (https://www.hdfgroup.org). The latter format has significant advantages in terms of file size (typically 10% of equivalent in XML) and speed of reading/writing.

**Supplementary Fig. 2** illustrates how these libraries can be used during the process of converting and sharing models on OSB. All of these libraries are installed and configured for use with supported simulators in the OSB Docker image (**Supp. Table 2**).

### NeuroML and PyNN

PyNN allows a single script in Python to be written which can instantiate and run a network in either NEURON, NEST, Brian or on neuromorphic hardware^55^; the only difference in the script is in the import of the package specific for that simulator (from *pyNN.neuron import* *, *from pyNN.nest import* *, etc.)^31^. The NeuroML export from PyNN works similarly (*from pyNN.neuroml import* *) and will export the cell parameters, connectivity and inputs to valid NeuroML. An example of this is the (increasingly widely used^56–58^) cortical column model^9^ in **Fig. 4e-h**. The original conversion of this model to PyNN was extended with 3D distributions for cell populations without changing the overall network behavior, but making the structure of the network clearer when visualized on OSB.

### Executing simulations on OSB

Having models specified in NeuroML format on OSB means these can easily be converted to a number of simulator specific formats for execution (**Supplementary Table 1**). A user who is signed in on OSB and is viewing a NeuroML model can “persist” the 3D view (orange star on top icon bar, e.g. **Fig. 3a**), so that that version of the NeuroML file (along with the layout/visual properties/current widgets) is stored for opening again at any time. Simulations can be set up and run with a number of options including simulation duration, time step, what to record (e.g. all membrane potentials at cell somas), and which simulator to run it on. Currently supported simulators include NEURON, NetPyNE, jNeuroML. The OSB website is hosted on an Amazon Web Services (AWS, https://aws.amazon.com) instance, on which the supported simulators are installed. Short simulations can quickly be run on a single processor on this AWS platform. Users are presented with a list of running and completed simulations (**Fig. 5**) and can plot recorded values, or overlay them on cells for visualization of network dynamics.

An alternative option for running simulations is enabled through integration with the Neuroscience Gateway^49^ (NSG, **Fig. 1a**). OSB generates the simulator scripts and submits these through the RESTful API of NSG (NSG-R, http://www.nsgportal.org/guide.html). These are then submitted to the Extreme Science and Engineering Discovery Environment (XSEDE) high performance computing resources for execution, where NSG has pre-configured multiple neuronal simulation packages. Users on OSB do not require any account on NSG or XSEDE, and completed simulations appear in the OSB interface in the same way, albeit with more of a delay than running on OSB’s own servers. Because NSG provides access to parallel computing, OSB users can select NetPyNE^38^ (http://netpyne.org) as a target simulator, and opt to run network models on up to 256 processors, significantly speeding up the simulation time. This number can be increased in future as we gauge usage statistics/demand, to make greater use of the thousands of processors available via NSG. Scripts have also been developed which allow modelers to submit NeuroML and PyNN directly to NSG-R, bypassing the web interface (https://github.com/OpenSourceBrain/NSGPortalShowcase).

Data generated during simulations launched via OSB can be downloaded from the web interface, but can also be set to automatically be added to the user’s Dropbox (https://www.dropbox.com) folder, to enable local analysis of the simulation data. This is enabled by generating an API key on the Dropbox site and adding this to the user’s OSB account.

Existing tutorials for OSB are listed at http://www.opensourcebrain.org/tutorials. Documentation for those interested in developing and hosting tutorials and making use of the visualization and simulation features of OSB can be found at http://opensourcebrain.org/docs#Creating_Tutorials.

### Converting neuroinformatics resources

The cell models used in the Blue Brain Project rat somatosensory microcircuitry network^8^ have been made available in the original NEURON format on the Neocortical Microcircuit Collaboration Portal (NMCP)^59^. We have developed scripts for automatically converting these cells to NeuroML and examples have been made available on http://www.opensourcebrain.org/projects/blue-brain-project-showcase. We have also converted the ion channel models from the Channelpedia^60^ database to NeuroML format in this repository. Representative connectomes with point neurons used in recent studies of the Blue Brain Project microcircuit^61, 62^ are available on the NMCP and a NeuroML HDF5 based version of this full connectome can be found on the OSB project. The full network (31346 cells, 7.6 million connections) cannot yet be displayed on OSB (though this is a target for future releases). However a scaled down version, with 5% of neurons in the original is available and can be visualized.

The Allen Cell Types Database^19^ (http://celltypes.brain-map.org) contains neuronal reconstructions and electrophysiological recordings from multiple cells in mouse visual cortex. The project has also made computational models of these cells available on the website in both biophysically and morphologically detailed (implemented in NEURON simulator) and point neuron (Generalized Linear Integrate and Fire (GLIF) models in a custom Python simulator) formats. These have both been converted to NeuroML/LEMS formats on http://www.opensourcebrain.org/projects/alleninstituteneuroml. Each cell shown in **Fig 3a** has a unique complement of ionic conductance densities tuned to the original cell’s electrophysiological recordings.

Neuronal reconstructions from NeuroMorpho.Org^63^ are available in standardized SWC format^51^ and these files, containing information on the 3D locations, radii, connectivity and type of the points in the reconstruction, can be visualized directly on OSB by placing them into a GitHub repository (see http://www.opensourcebrain.org/projects/neuromorpho). It is also possible to convert these files to NeuroML, to build them into spiking neuron models with active conductances, by using the application at https://github.com/pgleeson/Cvapp-NeuroMorpho.org.

The Janelia MouseLight project provides SWC versions of their neuronal reconstructions, in addition to a proprietary JSON file format with extra metadata. The OSB project for this (http://www.opensourcebrain.org/projects/mouselightshowcase) provides scripts for converting the JSON files to NeuroML while retaining the metadata, and the repository contains example converted files for visualization on OSB.

### Testing and model validation

The OSB Model Validation framework (OMV, https://github.com/OpenSourceBrain/osb-model-validation) was developed to facilitate testing and comparing model behavior across different simulators and formats. Expected behaviors (e.g. spike times, resting membrane potential) are described in short Model Emergent Properties (mep) files, as well as consistency checks for parameters such as total membrane area and temperature. Then a number of OSB Model Test (omt) files are written for each of the simulator configurations (engines) which should produce that behavior, specifying the allowed tolerance. Simulator entries in the last 3 columns of **Supp. Table 1**. are generally associated with individual passing tests (one omt file per simulator) linked by producing the same behavior as described in a shared mep file.

OMV can be installed locally and run at command line (see **Supp. Table 2**) and is also used on the continuous integration^64^ service Travis-CI (https://travis-ci.org). Each time there is a commit to a GitHub repository for a model, OMV is launched during the test on Travis-CI, the appropriate simulators are installed and all of the OMV tests in that repository are set running. This facilitates finding any changes to the model which break its intended behavior.

### Code availability

All of the code for the OSB platform and the models presented here is open source. The Ruby on Rails frontend for OSB is at https://github.com/OpenSourceBrain/redmine and the Geppetto repositories are listed at https://github.com/openworm/org.geppetto. The majority of repositories for the models in **Fig. 2** can be found at https://github.com/OpenSourceBrain.

## Figure Legends

**Supplementary Figure 1.**
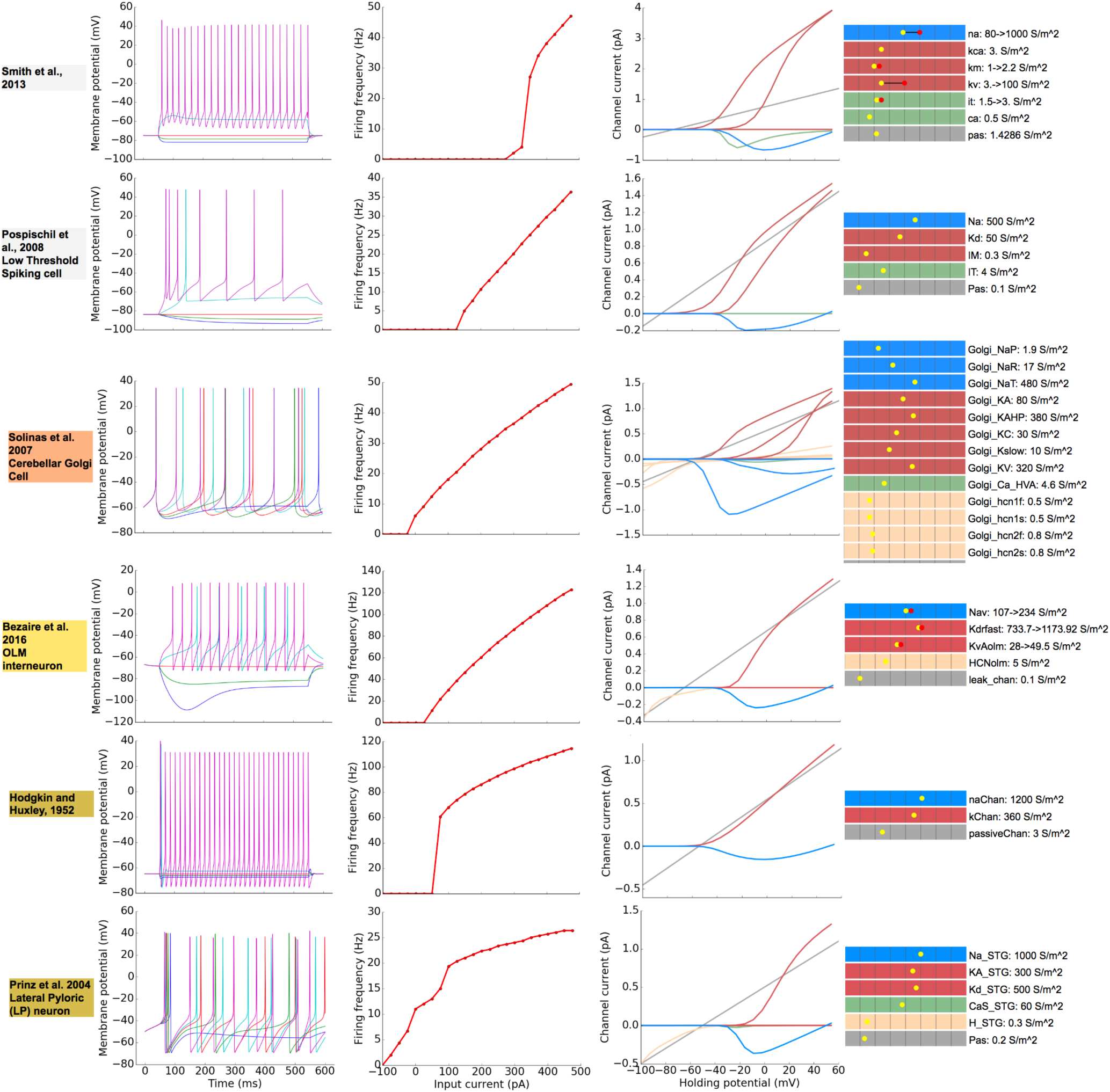
Comparison of the properties of cells and membrane conductances on Open Source Brain. Once neuronal models have been converted to NeuroML format their physiological properties can be readily compared, using same scripts to simulate and analyze the properties of different models (e.g. in Python using libNeuroML (Methods)). From left to right the columns show: the time course of the somatic membrane potential for 5 levels of somatic current injection (column 1); firing frequency as a function of current injected (column 2); currents through each conductance present in the soma when the cell is clamped at various holding potentials for the same conductance value (for the default minimal conductance of 10 pS in this case; column 3, colors correspond to colored scale in the next column); densities of the ionic conductances (column 4; logarithmic color scale shows densities of conductances grouped according to the ions transmitted: blue-Na^+^, red - K^+^, green - Ca^2+^, light brown - non selective HCN channels, grey - passive conductances; yellow dot is value if conductance is uniform, yellow is minimum and red maximum if nonuniform).

**Supplementary Figure 2.**
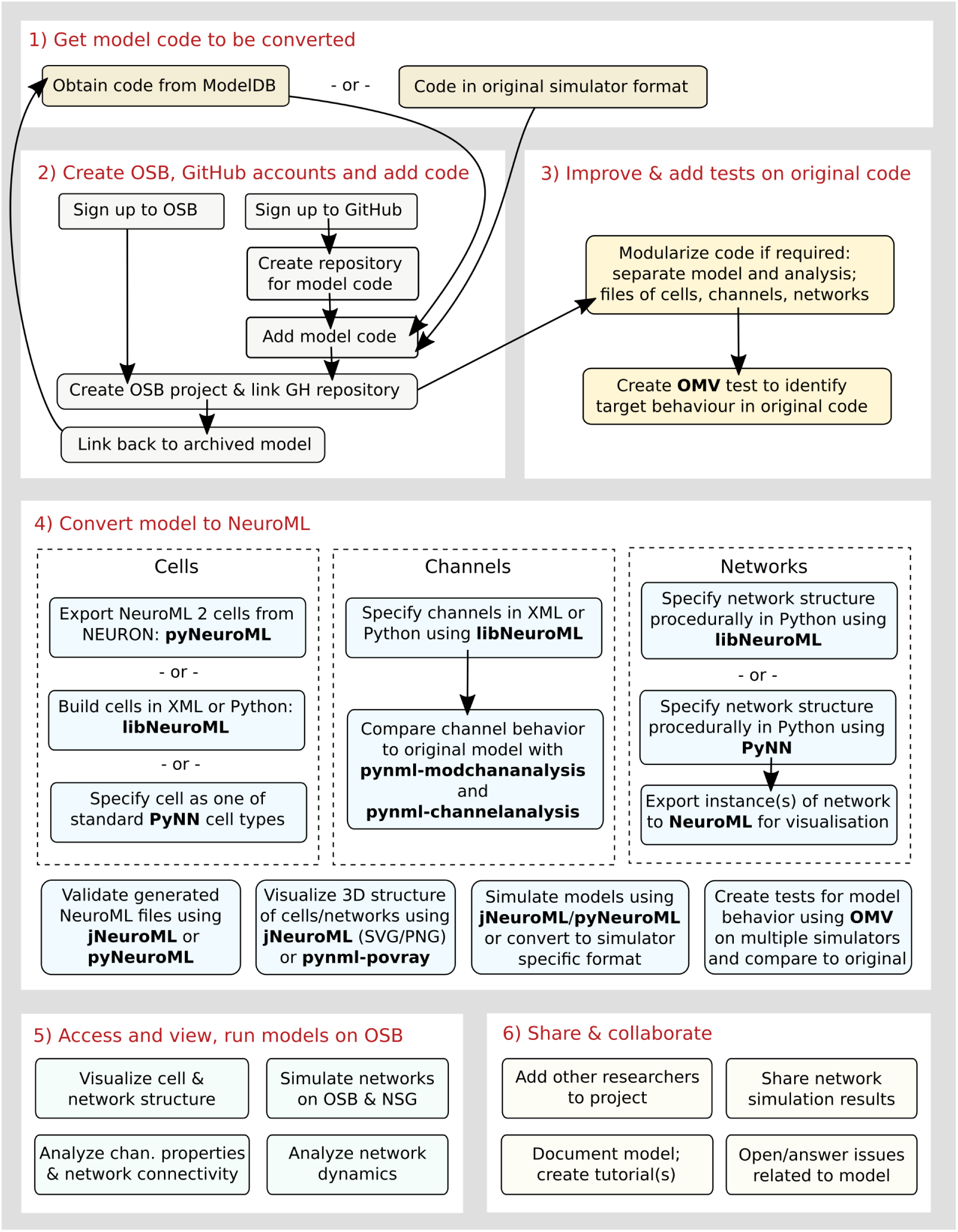
Procedures and tools to convert models from native formats into NeuroML and PyNN on Open Source Brain. Outline of the steps required for converting a model from its native language into NeuroML/PyNN format in order to utilize the visualization and simulation functionality on Open Source Brain (OSB) and for utilizing automated validation and other collaborative development tools. 1) The model code in the original format from the user or obtained from repositories such as ModelDB^32^. 2) User accounts are created on OSB and on GitHub. A new repository is created on GitHub and linked to an OSB project. 3) Optionally, the code should be cleaned up, further documented, and made more modularized, with all changes recorded in the version control system on GitHub. OSB Model Validation framework (OMV) tests can be added to the scripts which record the behavior of cell(s) in the original model and that the converted model needs to replicate. 4) The numerous tools available to build, export (e.g. from NEURON), validate, visualize and simulate the NeuroML/PyNN version of the model elements on a user’s local machine. As the structure of the model is replicated in NeuroML/PyNN, its properties can be tested against the behavior of the original model using OMV. 5) Once valid model elements are uploaded to GitHub, these can be visualized, analyzed and simulated on OSB. 6) Options available to make the model accessible, to get input from other users on the model and to share development responsibilities.

**Supplementary Table 1.** Automated testing of models on Open Source Brain. The models shown in **Fig. 2** have been tested for the validity of their NeuroML implementation and against expected behavior across a number of simulators. Colors in the first column are those used in **Fig. 2**. Tests include: (column 2) validity of the NeuroML files in the repository; (column 3) behavior of the PyNN scripts on different simulator backends (e.g. Brunel 2000 can be run in Brian, NEURON and Nest and exported to NeuroML), or conversion of the NeuroML files to PyNN scripts and execution on a simulator (e.g. NEURON for Ferguson et al., 2013); (column 4) execution of the NeuroML models using jNeuroML’s own simulator (point neuron models only); (column 5) execution of the NeuroML models in NEURON, translated using jNeuroML; (column 6) execution of native simulator code in the repository (e.g. original NEURON, Brian or Octave code taken from ModelDB) or other simulator code (NetPyNE, Brian 2) produced by converting the NeuroML representation using jNeuroML. Note, some models are not fully valid NeuroML because they have custom LEMS components. The current testing status of these and all other projects on OSB can be found here: http://www.opensourcebrain.org/status

| Main publication | Valid NeuroML | PyNN | jNeuroML | jNeuroML to NEURON | Other tests |
| --- | --- | --- | --- | --- | --- |
| Allen Institute Cell Types DB (Hawrylycz et al. (2016)) | ✓ |  | ✓ | ✓ | NEURON |
| Brunel (2000) | ✓ | Brian; NEURON; Nest; NeuroML | ✓ | ✓ | PyNEST; PyNN→ NeuroML |
| Hay et al. (2011) | ✓ |  | ✓ | ✓ |  |
| Izhikevich (2003) | ✓ | Brian; NEURON; Nest | ✓ | ✓ | Octave; PyNEURON |
| Markram et al. (2015) | ✓ |  | ✓ | ✓ | NEURON |
| Pospischil et al. (2008) | ✓ |  | ✓ | ✓ | NEURON; NeuroML→ Moose |
| Potjans and Diesmann (2014) | ✓ | Nest |  |  | NEST; PyNEST |
| Dura-Bernal et al. (2017) | ✓ |  | ✓ | ✓ | NeuroML→ NetPyNE |
| Smith et al. (2013) | ✓ |  | ✓ | ✓ | PyNEURON; NeuroML→ NetPyNE |
| Traub et al. (2005) | ✓ |  | ✓ | ✓ | neuroConstruct; NeuroML→ NetPyNE |
| Cayco-Gajic et al. (2017) |  |  | ✓ |  |  |
| Maex and De Schutter (1998) | ✓ |  | ✓ | ✓ | NeuroML→ NetPyNE |
| Solinas et al. (2007) | ✓ |  | ✓ | ✓ | NEURON |
| Vervaeke et al. (2010) | ✓ |  |  | ✓ |  |
| Bezaire et al. (2016) | ✓ |  | ✓ | ✓ | NEURON; NeuroML→ NetPyNE |
| Ferguson et al. (2013) |  | NeuroML→ PyNN→ NEURON | ✓ | ✓ | Brian; NeuroML→ Brian2 |
| Migliore et al. (2005) | ✓ |  | ✓ | ✓ |  |
| Pinsky and Rinzel (1994) |  |  | ✓ |  |  |
| Wang and Buzsaki (1996) |  |  | ✓ | ✓ | Brian; NEURON |
| Migliore et al. (2014) | ✓ |  |  | ✓ | PyNEURON |
| Boyle and Cohen (2008) | ✓ | NeuroML→ PyNN→ NEURON | ✓ | ✓ | Octave |
| FitzHugh (1961) | ✓ |  | ✓ | ✓ | NeuroML→ Brian; NeuroML→ Brian2 |
| Hodgkin and Huxley (1952) | ✓ | NeuroML→ PyNN→ NEURON | ✓ | ✓ | NeuroML→ Brian2; NeuroML→ Moose; NeuroML→ NetPyNE |
| Prinz et al. (2004) | ✓ |  | ✓ | ✓ |  |
| NeuroMorpho.Org | ✓ |  |  |  |  |
| Janelia MouseLight | ✓ |  |  |  |  |

**Supplementary Table 2.** Obtaining all core OSB models and testing against supported simulators. An image (preconfigured computational environment with files, libraries, etc.) for Docker has been created containing all of the simulators currently supported by OSB, along with all of the models presented in **Fig. 2** and **Table 1**. Once Docker is installed (on Windows, Mac OS or Linux), the image can be pulled from the central Docker registry. A container using this image can be set running, and OMV used to run tests on each of the models, in all the simulators it supports. The commands in blue above need to be typed at the command line. Alternatively, the Kitematic application bundled with Docker can be used for steps 2) and 3), searching for the image *opensourcebrain/simulation:osb_models-v0.7* and creating a container. The commands in 4) can then be entered into a command line terminal connected to this.

| Steps | Instructions |
| --- | --- |
| 1) Install Docker | Available from https://www.docker.com. |
| 2) Pull image with all simulators and stable projects | `docker pull opensourcebrain/simulation:osb_models-v0.7` |
| 3) Start a container with the image | `docker run -it opensourcebrain/simulation:osb_models-v0.7 /bin/bash` |
| 4) Inside container, go to folder with models and run OMV | `cd coreprojects`<br>`omv all` |

